# An actomyosin-mediated mechanical mechanism for brain neural tube elevation

**DOI:** 10.1101/2025.05.15.654327

**Authors:** Juana De La O, Chidera Okeke, Gabriel L. Galea, Adam C. Martin

**Affiliations:** Department of Biology, Massachusetts Institute of Technology, Cambridge, MA 02142, USA; Whitehead Institute of Biomedical Research, Cambridge, MA 02142, USA; Department of Developmental Biology and Cancer Researching and Teaching, University College London Great Ormond Street Institute of Child Health, WC1N 1 EH London, United Kingdom

## Abstract

Embryos fold their tissues into increasingly complicated shapes during development. Cells produce and coordinate the forces needed to fold tissues through networks of F-actin and Myosin II (actomyosin). We can discern the mechanical mechanisms used to fold tissues by analyzing these networks. Apical actomyosin is required to fold the neuroepithelium (NE), with a midline hinge and lateral neural folds into the neural tube. However, the large size and complex tissue curvature has complicated a detailed analysis of its actomyosin networks. Here, we developed a computational workflow to create 2D apical shell reconstructions of the NE. Using these projections, we confirmed a midline-lateral gradient of apical cell area and discovered a negative correlated gradient of actomyosin density. We hypothesized that lateral neural folds, with high apical cell constriction and actomyosin, have higher apical tension than the midline hinge, with large apical cell area and low actomyosin. Through target laser ablations in live embryos that allow us to infer tension, we confirmed that tension is isotropic and low at the midline and anisotropic and high on the lateral neural folds. Finally, we identified sex differences in cell shape, apical constriction rates, actomyosin in the NE at an earlier time point than previously appreciated. We use these findings to propose a lateral tension mechanism used in murine NTC that is distinct from the contractile hinge mechanism identified in other model systems.

## INTRODUCTION

As embryos develop, they continually sculpt their tissues into more complex three-dimensional shapes. For embryos to undergo such dramatic shape changes, the cells within and adjacent to a tissue must create and coordinate mechanical force with high spatial precision^1–3^. These cellular forces are in part generated by the cytoskeletal protein, filamentous actin (F-actin), coupled to the motor protein, non-muscle myosin II (henceforth myosin II). The F-actin and myosin II, collectively termed actomyosin, inside an individual cell can be connected to actomyosin pools in adjacent cells through transmembrane junctions^4–6^. When actomyosin connects across hundreds or thousands of cells, tissue-wide supracellular networks are formed that allow for the generation and coordination of tissue reshaping forces^7^.

One such supracellular actomyosin network can be found on the apical surface of the folding neural tube (NT)^8–13^. NT closure (NTC) is a phased folding process^14,15^. In the first phase, the flat neuroepithelium (NE), bends and creates a midline hinge and distinct lateral neural folds. In the second phase, the lateral neural folds elevate dorsally. In the final phase, the distal tips of the neural folds bend towards each other, appose, and fuse to form a sealed NT. The NT subsequently develops into the brain and spinal cord. While supracellular actomyosin networks can be found on the apical surface of the whole tissue, not all regions of the NE are equally reliant on this network to fold; Perturbations of actomyosin specifically lead to NT defects (NTDs) in the brain NT^16^. The brain NE is therefore an ideal system to interrogate mechanisms of actomyosin based tissue folding.

There is a menagerie of tissue shapes across developmental systems, so one might expect to see equally diverse applications of supracellular actomyosin networks. Instead, a core set of folding mechanisms are repurposed and adjusted to create a multitude of unique shapes and folds. For example, frog and chicken NTC uses a variation of the contractile hinge mechanism also observed in invertebrate tissue folding^17–19^. In this mechanism, cells at the future midline hinge point are enriched with apical actomyosin. Cells at the midline then constrict their apical surface and transform from a column to a wedge shape^20,21^. The regional constriction of cells into wedges creates a hinge in the tissue. This hinge can be propagated along the length of the tissue to form a crease. In chicken NTC, cells constrict anisotropically, with the greatest constriction orthogonal to the length of the crease^21^. In frog NTC, isotropically constricting cells are interspersed with anisotropically elongating cells^22–24^. In both cases, active actomyosin-mediated contractility at the hinge is proposed to be a primary driver for folding of the tissue.

At first glance, one might expect the mammalian NT to also employ the contractile hinge mechanism. In terms of broad shapes, the murine brain NE also develops a midline hinge and its closure is also diminished by actomyosin perturbations^10,12^. However, looking beyond superficial similarities, we find there are key discrepancies. While actomyosin perturbations do prevent full brain NTC, they do not disrupt formation of the midline hinge^9,25–27^, suggesting the hinge forms through an actomyosin independent mechanism. Consistent with this, cells at the midline hinge do not apically constrict, but the cells in the lateral neural folds do^28^. Additionally, unlike other organisms, the murine NT faces unique morphological constraints that must be overcome to complete folding^29^. First, in the initial phase of NTC the lateral neural folds have pronounced dorsal bulges that interfere with elevation and apposition unless their curvature is inverted. Second, the murine NT has substantial global curvature from a rostral-caudal flexure underlying the brain region. The contractile hinge mechanism of bending is insufficient to explain these midline behaviors and does not provide a means to resolve these additional morphological hurdles. What then are the actomyosin-mediated mechanical mechanisms employed during mouse brain NE elevation?

Here, we set out to answer this question. We started by developing a new approach to visualize and quantitatively analyze apical actomyosin networks on the large, curved surfaces of the NE. Using this framework, we reevaluated the medial-lateral pattern of apical cell constriction and discovered an inversely correlated pattern of actomyosin intensity. We predicted that this regional pattern of actomyosin-mediated apical constriction leads to a regionalized pattern of tissue tension, such that the lateral neural folds exhibit higher tension than the midline hinge. We confirmed this through a series of targeted laser ablations in live mouse embryos, that allowed us to measure recoil as a proxy for tissue tension. We also found that the directional bias in cell shape and orientation was predictive of tension anisotropy in different regions of the NE. Finally, throughout our experiments, we found differences in cell shape, constriction rates, and tissue mechanics between female and male embryos. Our work defines a lateral tension mechanism of actomyosin mediated tissue folding and provides some of the first mechanical and cellular evidence for sex differences during mouse NTC.

## RESULTS

### An imaging processing workflow to analyze apical actomyosin networks of neural tubes

Work in other systems has shown that studying how actomyosin networks are composed and structured can inform on the mechanics and mechanisms of tissue folding^18,22,30–35^. However, the curious morphological constraints of the murine brain NE present a challenge for visualizing and analyzing the apical actomyosin networks. The tissue curvature makes it so that no single imaging plane captures all the apical information in a field of view. The large size of the brain NE, on the order of 500 µm wide and 800 µm long (Figure 2B and C), also typically requires a trade-off between whole tissue or subcellular resolution images. This is particularly encumbering as the murine NT does not close uniformly, rather progressively zippering close from discrete closure points^14–16^. As such, mechanical constraints, and by proxy the structure of the actomyosin network, are likely to vary along both the midline-lateral and the rostral-caudal axis. Therefore, to accurately measure and compare the actomyosin networks across samples and states of development, we need to connect cell-level behaviors to the location of cells within the folding NE.

We accomplished this by developing an imaging and analysis workflow that allows us to project a superficial image of the NE with sub-cellular resolution. We started by fixing, staining, and mounting whole embryos between glass coverslips. We then tile-scanned the dorsal face of the midbrain and hindbrain NE, ensuring regions of overlap between field of view (Figure 1A). Through a combination of FIJI macros, Ilastik, and custom written MATLAB script, we extracted a “shell” of only the most apical 4.5 µm of information in each field of view (Figure 1B). We then stitched together individual shell projections to reconstruct the whole dorsal NE surface (Figure 1C). We segmented these reconstructions using Cellpose^36^ (Figure 1D) to extract key measurements (Figure 1E).

**Figure 1:**
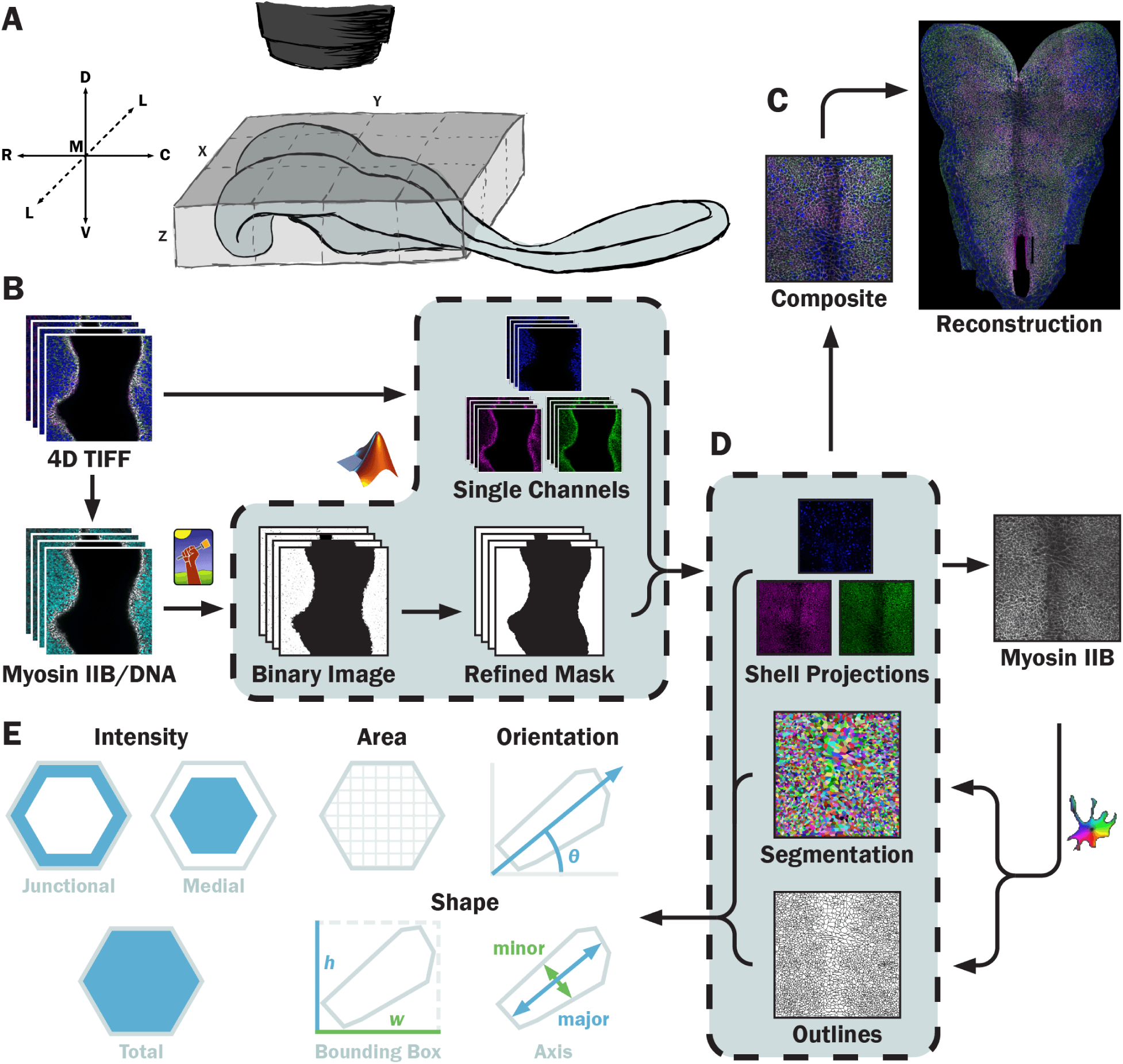
An imaging processing workflow to analyze apical actomyosin networks of neural tubes. **A**) A schematic of tile scanning whole fixed mouse embryos. We place samples between glass coverslips and scan en face, with the Z imaging axis parallel to the dorsal-ventral axis. **B**) The resulting confocal stacks are then processed through our computational workflow. A Myosin IIB and DNA composite is inputted to Ilastik, which bins pixels as part of the sample or background. The binary image is refined into a mask which is used to create single-channel projections (see Methods). **C**) Projection composites are computationally stitched together, reconstructing the brain neural tube and allowing for tissue-level measurement. **D**) Cellpose uses the Myosin IIB projection to segment and outline cells, enabling key cell measurements (**E**). Shaded boxes in B) and D) indicate processed completed by custom MATLAB scripts. Dorsal (D), ventral (V), rostral (R), caudal (C), lateral (L), medial (M). X, Y, and Z denote image axes.

### A midline-lateral gradients of apical constriction in the midbrain vary between sexes

Armed with a new visualization framework, we began by investigating the patterns of apical constriction. A midline-lateral gradient of apical constriction was previously reported in the midbrain^28^. Cells in the midbrain start off large at earlier stages, but as NTC proceeds, cells on the lateral neural folds constrict while midline hinge cells do not significantly change their apical area^28^. We replicated these results with a few key adjustments.

First, we binned embryos strictly by their somite stage (ss) rather than pool embryos in adjacent stages together. Somites — transient masses of cells that form in the embryo — appear in pairs every 1-2 hours starting at Embryonic Day (E) 8, allowing for higher precision stage-matching of embryos. Because the broad shapes of the NE changes rapidly and predictably between stages^29^, the corresponding cell shapes and actomyosin networks are also likely to change from one stage to the next.

Second, we standardized cell position along the midline-lateral (ML) axis as a percentage distance from midline rather than microns. The murine brain NE changes width throughout NTC, progressively widens until 5-6 ss, after which it narrows as the neural folds elevate and begin to appose^29^. Additionally, variance in embryo mounting may cause opposite neural folds to have different apparent widths within the same embryo. By plotting cell position as a percentage, we corrected for differences in tissue width between stages and variation due to embryo mounting, allowing us to compare equivalent positions.

We focused our attention on the midbrain region of the NE during the elevation phases, of 5-8 ss as younger embryos have midlines that are too sharply bent to analyze. The dorsal bulges of the neural folds also begin to flip curvature at these stages^29^. We analyzed all the cells in a 100 µm tall band across the midbrain of embryos (Figure 2A-C). This zone sits rostral to and excludes the cells that compose the midbrain/hindbrain boundary, which have large midline-laterally elongated cell morphologies that would skew our measurements (Figure 2A-C). Consistent with prior literature^28^, we find that at 5 ss, cells are uniformly large along the ML axis. However, we also find that males consistently have larger cells than females (Figure 2D, F, and H). By 8 ss, both males and females have larger apical areas at the midline (10-20% distance) and the boundary of the NT (90-100% distance) and smaller apical areas on the lateral neural folds (30-80% distance) (Figure 2E, G, and I). However, at this stage females have significantly larger cell areas at the midline than males while males have larger cell areas at the NE-surface ectoderm (NESE) boundary.

**Figure 2:**
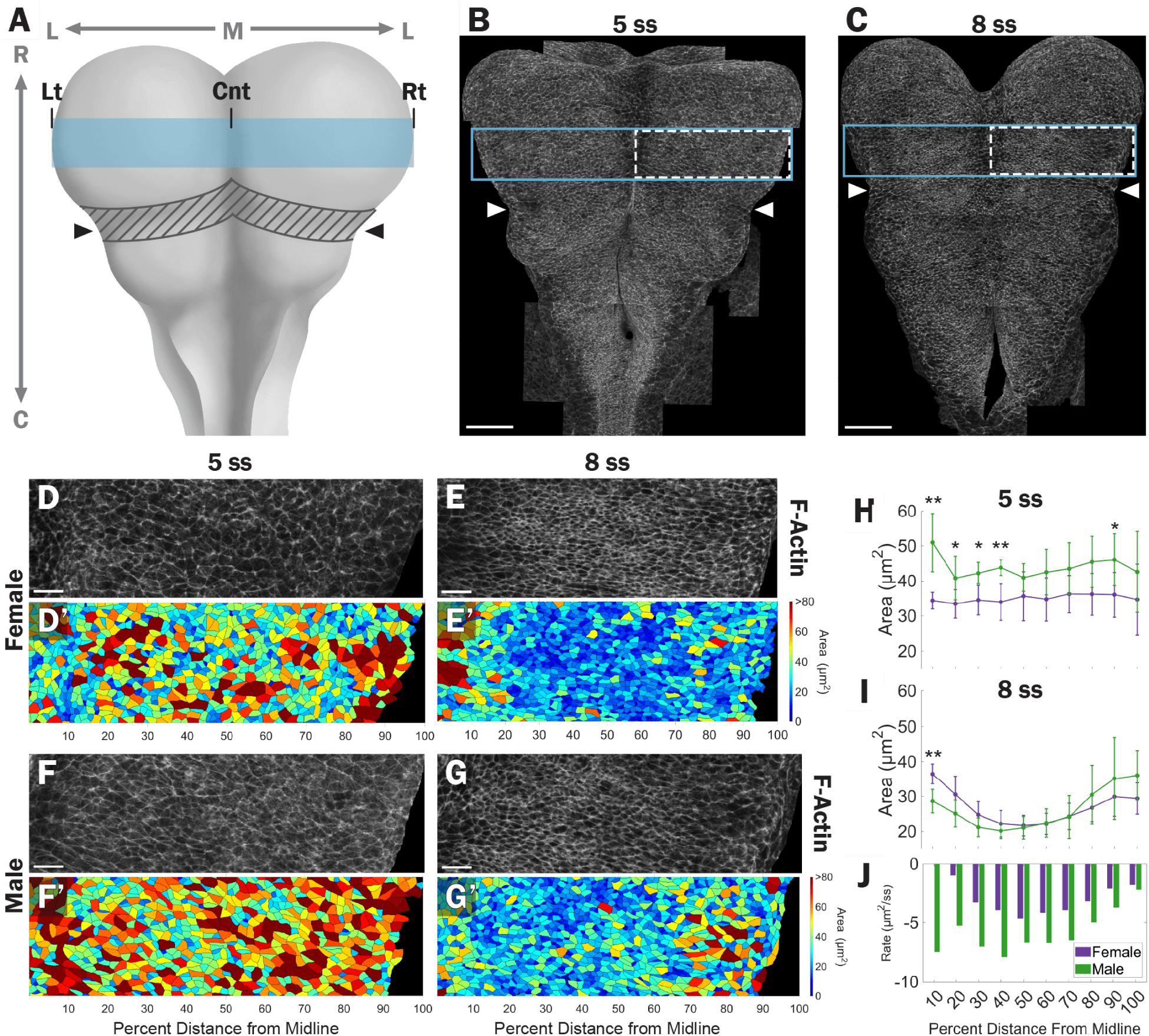
A midline-lateral gradients of apical constriction in the midbrain vary between sexes. **A**) A dorsal view cartoon of the mid- and hindbrain. Hashed grey area marks boundary cells that are excluded from analysis. Ticks indicate where the left, center, and right boundaries are called. **B** and **C**) Representative images of B) 5 and C) 8 ss embryos F-actin. A-C) Arrowheads denote the midbrain/hindbrain boundary. Blue boxes mark 100 µm tall region analyzed. Dashed white boxes indicate area where representative images are taken from. **D -G**) Representative F-actin images of D-E) female and F-G) male midbrains at D, F) 5 and E, G) 8 ss. D’-G’) Segmented images color coded by area from small (dark blue) to large (dark red). **H-I**) Plots of average cell area (dots) and standard deviation (bars) as a function of distance from midline at H) 5 and I) 8 ss. **J**) Rates of apical construction from 5 to 8 ss as a function of distance from midline. H-J) Females are purple and males are green. Rostral (R), caudal (C), lateral (L), midline (M), Left (Lt), Center (Cnt), Right (Rt). * p < 0.05, ** p < 0.01.

The number of cells across the NE does not change over this time^29^, suggesting that changes in apical area are not the result of cell division. Because the apical cell areas in males starts off larger than in females at 5 ss, for them to become smaller or equivalent in size to females by 8 ss, cells in males must constrict at a faster rate than in females. Indeed, when we calculate the average rate of apical constriction, we find it is greater in males than in females throughout the ML axis, except at the NESE boundary (Figure 2J). Despite these differences, both sexes exhibit the same regionalized cell behaviors: large apical area at the midline hinge and cells on the lateral neural folds. But what is driving these behaviors?

### A midline-lateral gradient of apical actomyosin negatively correlates to cell are in the midbrain

One possible driver of the regional patterns of apical constriction the apical network of F-actin and myosin IIB, a dominant myosin II isoform expressed in the murine NE^37,38^. Apical enrichment of actomyosin is known to facilitate selective constriction of cells^7,39,40^ and we predicted that cells that undergo greater apical constriction could be enriched in actomyosin relative to cells that do not. However, cell shape does not always match myosin contractility levels, because cell shape depends on extrinsic, as well as intrinsic forces^41^. We found that at 5 ss the large cells at the midline have lower amounts of both F-actin and myosin IIB density than cells on the lateral neural folds or NT boundary in both sexes (Figure 3A, C, E, G and Figure 3 – Supplement 1 A, C). Interestingly, females have a significantly higher density of myosin IIB than males at the midline (Figure 3G). This broad pattern of low midline and high lateral actomyosin density persists at 8 ss, with a slight increase in the F-actin density on the neural folds in both sexes, and an increase of myosin IIB density at the male midline (Figure 3B, D, F, H, and Figure 3 – Supplement 1 B, D). Therefore, actomyosin density negatively correlates with apical cell area at all regions except the NT boundary of 8 ss embryos. This pattern of actomyosin further argues against the use of the contractile hinge mechanism in murine brain NTC.

**Figure 3:**
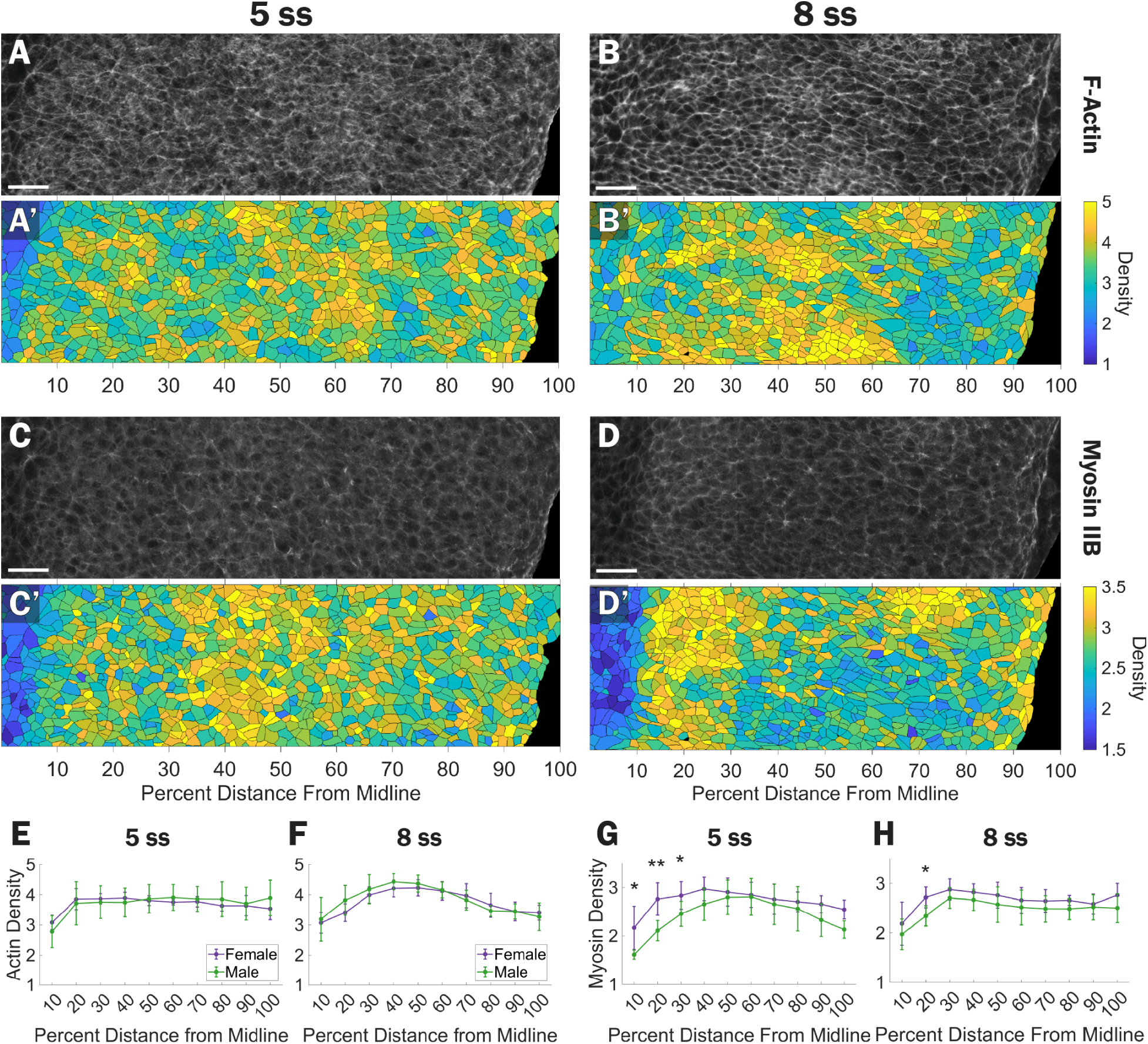
A midline-lateral gradient of apical actomyosin negatively correlates to cell area in the midbrain. **A-D**) Representative images of male midbrains stained for A-B) F-actin, and C-D) Myosin IIB at A, C) 5 and B, D) 8 ss. **A’-D’**) Segmented images color coded by density of protein expression from low (dark blue) to high (yellow). **E-H**) Plots of average cell E-F) F-actin and G-H) Myosin IIB density (dots) and standard deviation (bars) as a function of distance from midline at E, G) 5 and F, H) 8 ss. Females are purple and males are green. * p < 0.05, ** p < 0.01.

### The lateral midbrain neural folds exhibit higher tension than the midline in a sex dependent manner

Although there are nuanced differences between sexes, they both generally exhibit similar patterns: 1) Apical cell areas at the center are larger than on the neural folds, 2) substantial apical constriction occurs on the lateral neural folds, and 3) actomyosin density is lower at the midline and higher on the neural folds. We theorized that this might reflect regional differences in apical tissue tension across the midbrain, with the midline exhibiting less tension than lateral neural folds.

We put this theory to the test through targeted two-photon laser-ablations in the midbrain of live embryos^42^ (Figure 4A). We measured the velocity of recoil after ablation as a proxy for tissue tension; the faster the recoil orthogonal to the ablation, the greater the tension was in the tissue prior to ablation. Recoil velocity is influenced by the tissue viscosity and elasticity, dampening over time^43^. Therefore, we measured the initial recoil velocity (IRV), defined as the recoil velocity from the time of the ablation to the next immediate frame. We tracked points across the center of the ablation path to calculate IRV (Figure 4B). We also tracked points orthogonal and parallel to the direction of recoil to control for artifactual background movement, primarily from the embryonic heartbeat. Our recoil measurements averaged ∼2.8 µm/s, while both controls averaged close to 0 µm/s (Figure 4C). We conclude that our assay has sufficient sensitivity to measure tissue recoils that represent a release in pre-existing tissue tension.

**Figure 4:**
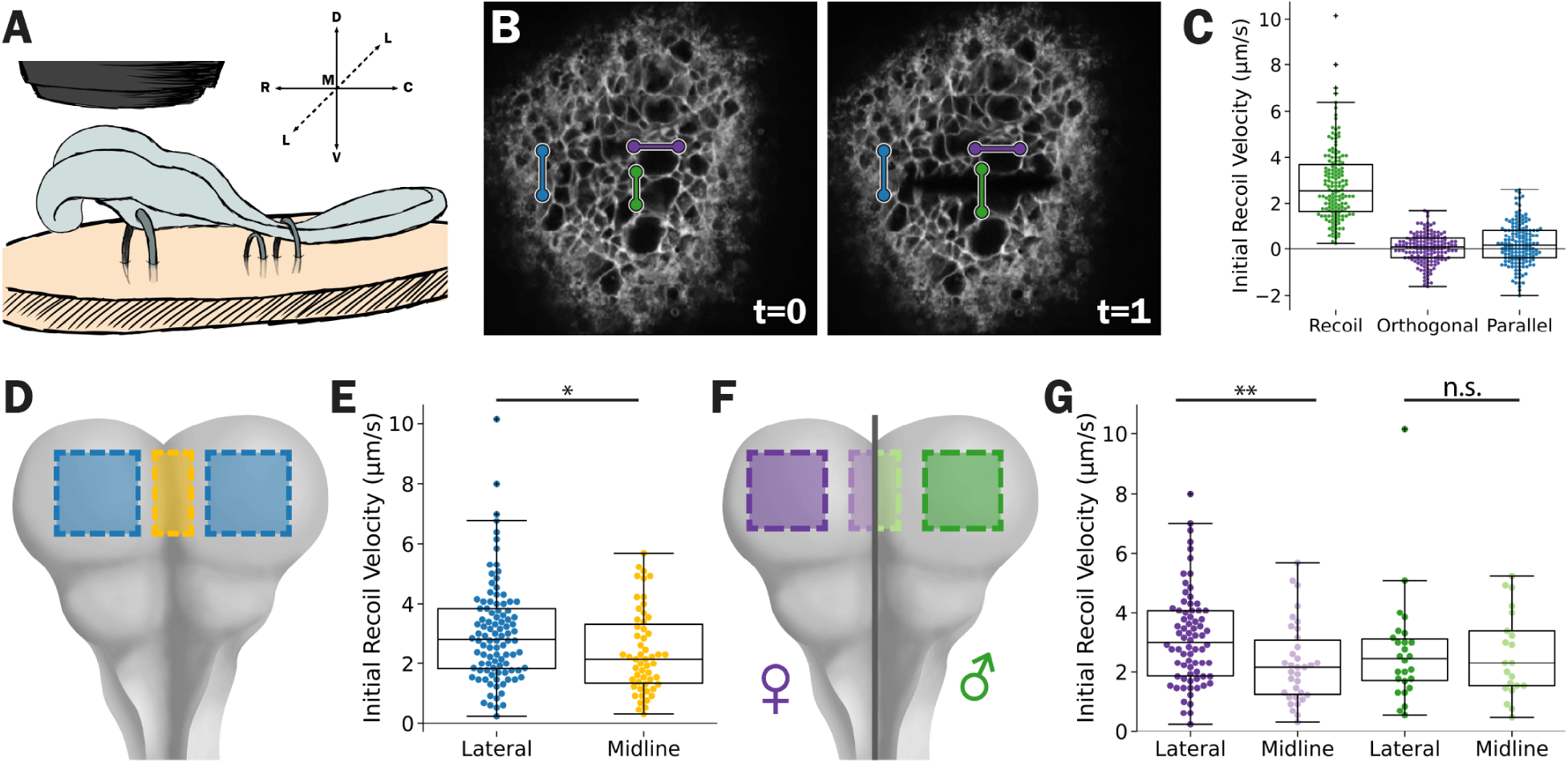
The lateral midbrain neural folds exhibit higher tension than the midline in a sex dependent manner. **A**) A schematic of live mouse embryo laser ablations. After partial dissection of support membranes and staining, embryos are positioned with microneedles embedded in an agarose plate. A dipping objective is used to ablate samples immersed in culturing media. **B**) Representative images of a CellMask stained embryo before (t=0) and immediately after ablation (t=1). Two points on opposing sides (green) of the ablation were tracked and used to calculate the initial recoil velocity (**C**). Points orthogonal (purple), and parallel (blue) to the ablation serve as internal controls. **D** and **F**) A dorsal view cartoon of the mid- and hindbrain. Shaded areas indicate regions where initial recoil measurements plotted in **E**) and **G**) where measured. In F-G) female measurements are in purple hues and males are in green hues. In E and G), box plots show the median and interquartile range. Dorsal (D), ventral (V), rostral (R), caudal (C), lateral (L), midline (M), midline-lateral (ML), rostral-caudal (RC). Not significant (n.s), * p < 0.05, ** p < 0.01.

We targeted our ablations to the midline (within 10% distance of the neural fold length) and lateral neural folds (∼70 % distance). Consistent with our predictions, the lateral neural folds have higher IRV (3.03 µm/s) than the midline (2.40 µm/s) on average (Figure 4D and E). When we disaggregate the data by sex, we found that females have a significantly higher IRV laterally than at the midline (3.15 µm/s vs. 2.30 µm/s, 37% difference). Strikingly, we did not detect a significant difference these regions in males, which showed a midrange of IRV at the midline and laterally (2.68 µm/s vs. 2,57 µm/s, 4% difference, Figure 4F and G). For all our measurements, we found no difference in IRV over the somite stages analyzed (Figure 4 – Supplement 1). The patterns of apical cell area, constriction, and actomyosin enrichment we measured were predictive of the low midline tension in both sexes and the high lateral neural fold in females. However, despite being nearly identical as the females, these patterns were not consistent with tension measurements in the male neural folds, leading us believe that there were more nuances in the tissue tension.

### Apical tension is isotropic at the midline and anisotropic on the lateral neural folds of the midbrain

It seemed unlikely to us that there would be a difference in regional tension between sexes when their cellular shapes and actomyosin expression were so similar. Rather, we reasoned that it was more likely that we had not yet considered an additional variable that had impacted the results, namely the directionality of the tension. Therefore, we looked for evidence of tension anisotropy. To start, we calculated the average apical cell aspect ratio, defined as the ratio of the major axis length over the minor axis length. A ratio close to 1 indicates a uniformly round cell shape while ratio greater than one indicates an elongated shape. At both 5 and 8 ss cells throughout the ML axis have an average aspect ratio of ∼2 in both sexes, indicating that cells’ apical shape are, on average, twice as long on their major axis as their minor axis (Figure 5 A-B, E-F, and Figure 5 – Supplement 1 A-B).

**Figure 5:**
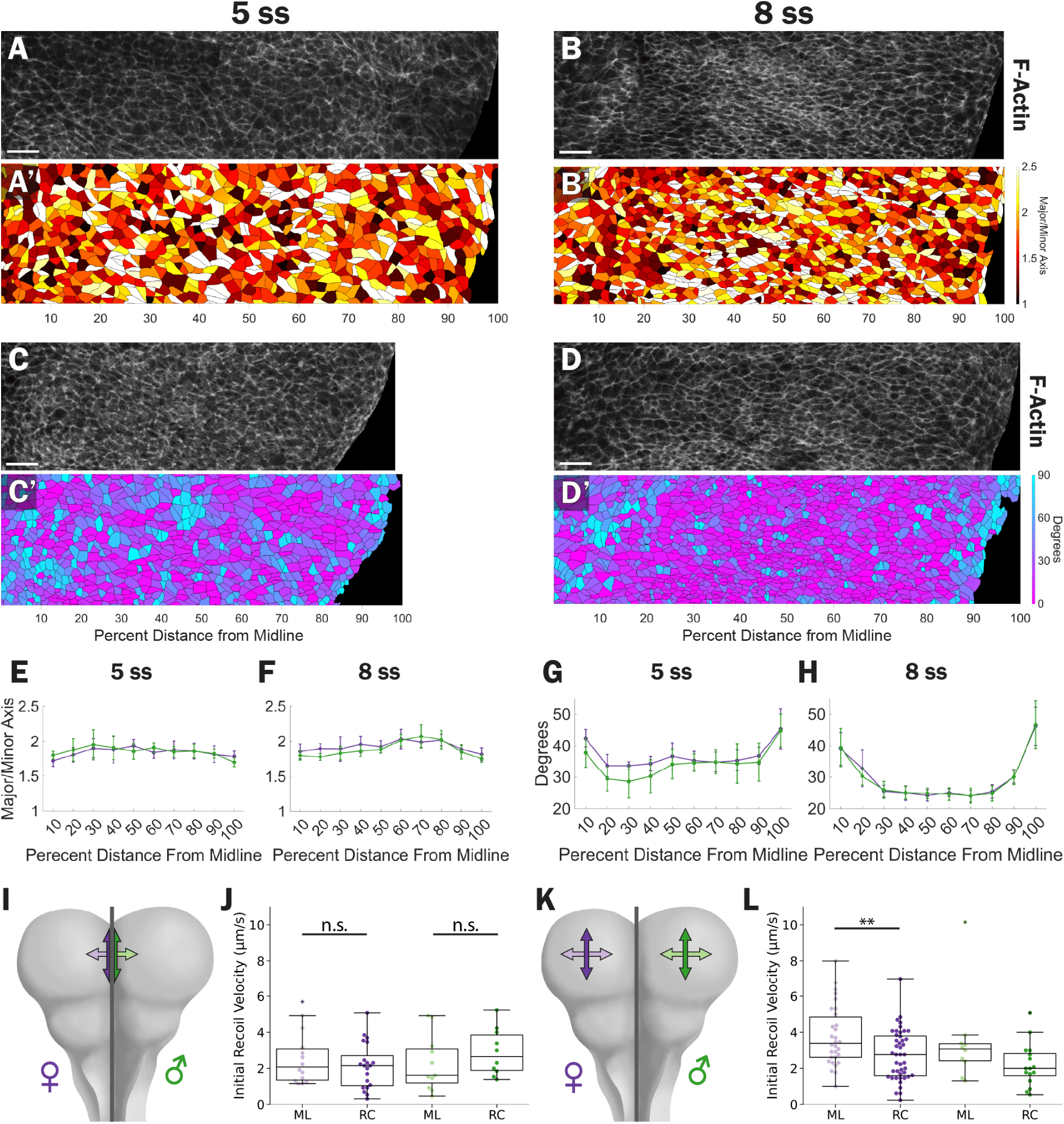
Apical tension is isotropic at the midline and anisotropic on the lateral neural folds in the midbrain. **A-D**) Representative F-actin images of female midbrains at A, C) 5 and B, D) 8 ss. **A’-B’**) Segmented images color coded by aspect ratio from uniformly round (black) to elongated (white). **C’-D’**) Segmented images color coded by degree from midline, with 0 (pink) being parallel to the midline-lateral axis and 90 (blue) being parallel to the rostral-caudal axis. **E-H**) Plots of average cell E-F) aspect ratio and G-H) orientation (dots) and standard deviation (bars) as a function of distance from midline at E and G) 5 and F and H) 8 ss. **I** and **K**) A dorsal view cartoon of the mid- and hindbrain. Double headed arrows indicate the direction inferred tension was measured in the **J**) midline and **L**) lateral neural folds. Box plots show the median and interquartile range. In E-H, I-L) female measurements are in purple hues and males are in green hues. Data shown is the same as Figure 4 but binned by orientation of initial recoil velocity. Not significant (n.s), * p < 0.05, ** p < 0.01.

We then determined the orientation of cells. Cells orientated to 0° are parallel to the ML axis while cells oriented to 90° are parallel to the rostral-caudal (RC) axis. At 5 ss, cells across the NE had an average orientation between 30° and 45°, indicating a minor ML bias in orientation (Figure 5C and G). By 8 ss, cells at the midline and NESE boundary maintained an average orientation of 40°-45°, indicating that although they are elongated, they have almost no directional bias. By contrast the cells on the lateral neural folds average ∼25°, indicating a stronger ML orientation bias (Figure 5D and H). These patterns were identical between both sexes (Figure 5 G-H and Figure 5 – Supplement 1 C-D). Our results suggest that at the midline and NESE boundary, though cells are elongated, they do not have directional bias and likely have isotropic tension. At the same time, the cells of the lateral neural folds, with an elongated shape and biased ML orientation, likely have higher ML tension than RC tension.

To test this prediction, we went back to our laser ablation data. As we performed ablations, we ensured that all ablations were oriented along either the ML or RC axis of the embryo. The resulting IRVs provide information on the RC and ML axis, respectively. For clarity, we will discuss direction of the measured IRVs and, therefore, inferred tension. As predicted by our cell shape analysis, IRVs are isotropic at the midline of both sexes (Figure 5I and J). On the neural folds, we found that females have higher ML than RC IRV (3.77 µm/s vs. 2.73 µm/s, 38% difference). We also found that males have higher ML than RC IRV (3.45 µm/s vs 2.19 µm/s, 56% difference, Figure 5K and L), although this did not reach statistical significance. For all our measurements, we found no difference in IRV over the somite stages analyzed (Figure 5 – Supplement 2). Given that the lateral neural folds of both sexes 1) undergo apical constriction, 2) are relatively enriched with actomyosin, and 3) have substantial cell shape anisotropy, we propose that the anisotropic tension is present in the neural folds for both sexes.

## DISCUSSION

We set out to define a mechanical mechanism for the elevation of the mouse brain NE. Using an image processing workflow we developed to create apical projections of the brain NE at subcellular resolution. In analyzing these images, we confirmed that cells on the lateral neural folds undergo apical constriction, while cells areas at the midline remain relatively large^28^. We discovered a ML gradient of F-actin and Myosin IIB density that negatively correlates with apical cell areas in later stage NTs, being high on the lateral neural folds and low at the midline hinge. Finally, and consistent with these results, we showed that the lateral neural folds have higher anisotropic tension relative to the midline hinge, which have lower isotropic tension. Combined with prior work showing that actomyosin perturbations do not prevent midline hinge formation^9,25–27^, our data argues against a contractile hinge mechanism being used during the elevation of the murine brain NE. But if not a contractile hinge, then what mechanism is used? How does this regional pattern of apical constriction and tension facilitate brain NT elevation?

### A lateral tension model for cranial neural tube closure

We propose that a lateral tension mechanism is used during elevation of the murine brain NE. In this mechanism, apical constriction in the lateral neural folds, mediated by high actomyosin, introduces high, anisotropic tension. Combined with tissue thickening^29^, the lateral neural folds become rigid and stiff^44^. At the same, the midline cells maintain large apical areas, a low density of actomyosin, and slightly thin over time^29^, making them more flexible and amenable to bending. In essence, the NE develops many of the same characteristics as a hard cover book (Figure 7A). This regionalized pattern of tissue tension and rigidity primes the NE to elevate in response to an external pushing and elevating force. The rigid lateral folds (the book covers) can efficiently elevate, while the flexible midline (the book spine) bends, creating the hinge (Figure 7B).

**Figure 6:**
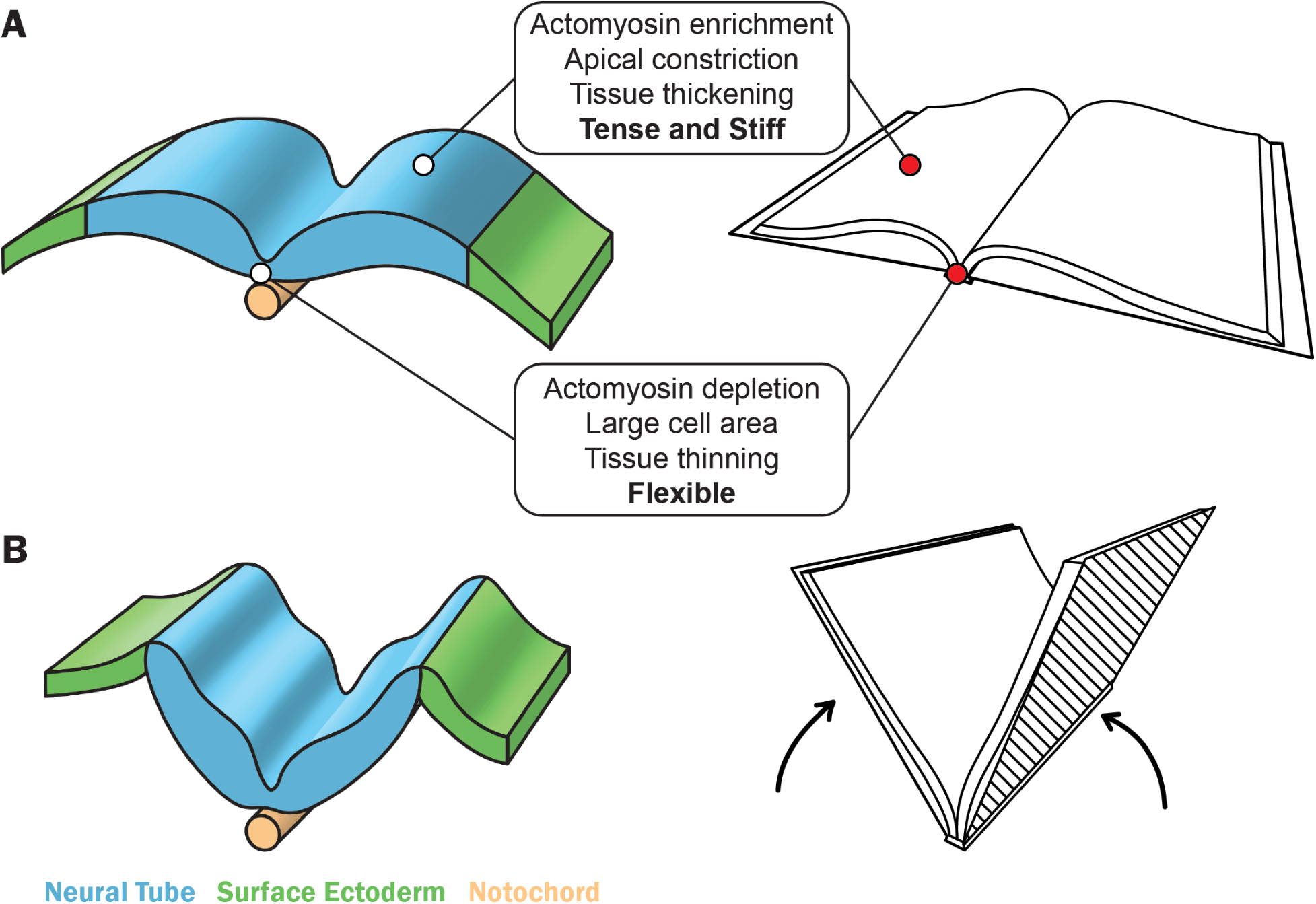
A lateral tension mechanism for brain neural tube elevation. Cartoons of the neural tube (left) and hardcover book (right) illustrating analogies between the two during **A**) early neural tube dorsal bulging and **B**) lateral neural fold elevation and midline bending. Arrows denote a hypothetical “pushing” force from adjacent tissues. Neural tube is blue, surface ectoderm is green, and the notochord is orange.

Our mechanism also provides a conceptual basis for understanding how the NT overcomes larger scale morphological challenges to elevate. Initially, the lateral neural folds bulge dorsally and exhibit convex curvature along the ML axis^29^. Left as they are these bulges would bump into each other as the neural folds elevate and prevent the apposition and fusion of the most lateral tips of the NT. The anisotropic tension in lateral neural folds and tissue thickening^29^ could contribute to the flattening and ultimate inversion of curvature to enable closure.

The formation of a hinge in the absence of actomyosin activity is also readily explained in this framework. At the midline hinge there is a low density of actomyosin and cells thin over time. Therefore, rather than bending from active actomyosin-mediated constriction, our results suggest that the midline hinge bends as a result from forces extrinsic to the NE that are generated independently from actomyosin. However, we cannot rule out that other tissue intrinsic cellular processes at or adjacent to the midline hinge contribute to tissue folding and hinge formation.

Our mechanism presupposes the existence of an external force “pushing” on the neural epithelium to promote elevation and midline bending. Where is this force coming from? We propose two likely, but not mutually exclusive sources. First is the mesenchyme, which underlies the NT, and is composed of migrating neural crest cells and cranial mesoderm cells. The extracellular matrix of the mesenchyme expands, and cells migrate concurrently with neural fold elevation and closure. When migration of cells within the mesenchyme or expansion of matrix is disrupted, NTC is prevented, suggesting to us that these process help promote tissue folding^45–47^. Second is the surface ectoderm, which is adjacent to either side of the NT boundary. Mutations specific to the surface ectoderm also result in NTC defects^48–50^. At the very least the surface ectoderm must relax or expand towards the midline to allow the neural folds to elevate and begin apposing. But we can also envision the surface ectoderm actively moving towards the midline and pushing in from either side of the NT, aiding in neural fold elevation and midline bending. You can observe a similar effect by folding a crease into paper and unfolding it, then placing your fingers at either end of the paper and pushing your fingers towards the center crease. Likely, a combination of multiple tissue intrinsic and extrinsic forces working in concert are required for NT elevation and final closure.

### Sex difference in early mouse embryo development

Past work has demonstrated that females have a higher penetrance of brain NTDS ^51^, implying that sex differences in the process must be present during or before NTC. It is nonetheless surprising that we have identified cell-level sex differences during NE elevation. Sex differences in embryo development have in part led to the low male samples in our laser ablation data. Male embryos are known to develop faster than female littermates^52,53^. At the time of sample collection more males had exceeded 10 ss, after which the neural folds are too elevated to ablate using our mounting procedures. As a result males were under-represented in our laser ablation data set (70% female, 30% male, Chi Square p = 0.0000830). We believe that both sexes demonstrate anisotropic tension, given that males have cell aspect ratios and orientations in the lateral neural folds as females. The averaging of RC and ML IRVs likely masked tension differences between the midline hinge and the lateral neural folds, creating the false impression that males did not have high tension on the lateral neural folds.

We have shown that females, despite having equivalent amounts of actomyosin as males on the lateral neural folds, constrict slower and yet paradoxically appear to have higher apical RC tension. One might expect that a slower contraction rate would introduce less tension into a tissue. However, tension is a complex measurement made of a balance of constricting and resisting forces from the perspective of the neural epithelium. A high tension and slow constriction rate, given equivalent amounts of actomyosin, suggests that there is an external high antagonistic force that is resisting constriction in females that is less prevalent in males. This is notably the same axis along which the brain NT ventrally flexes, a curvature and strain that must be overcome to complete NTC. A higher level of resistance and tension that females need to overcome in this direction could sensitize females to perturbations and increase the likelihood of developing an NTD.

If there are sex differences manifesting at the cellular and mechanical level within in the NT, where do they originate from? At this developmental age, the sex organs have not yet begun to develop, so a difference in hormonal inputs cannot explain these differences^54,55^. Instead, leading theories point to X-inactivation, which occurs before NTC, as the key genetic event that may drive a sex bias in NT defects^51^. How this chromatin-based event cascades into physical differences is not well understood, but a potential candidate is modulation of the Sonic Hedgehog (Shh) pathway, as changes in the expression of this pathway alter lateral neural fold apical constriction^28^. While not currently standard practice, there are strong arguments to factor sex as a biological variable in future studies of NTC and similar early developmental process.

## METHODS AND MATERIALS

### Animals

Mouse experiments using fixed samples described in this study were approved by the Massachusetts Institute of Technology Institutional Animal Care and Use Committee (IACUC) (institutional animal welfare assurance no. A-3125-01). We purchased FVB/NJ inbred mice (strain #: 001800 from Jackson Laboratories and bred in-house. We performed live mouse embryo studies under the United Kingdom Animals (Scientific Procedures) Act 1986 and the Medical Research Council’s Responsibility in the Use of Animals for Medical Research (1993). We obtained outbred CD-1 mice from the Charles River (CRADL, South Mimms, UK).

### Fixed Embryo Collection

We set up continuous mating pairs of FVB mice and checked every morning for the presence of a vaginal plug. The morning the plug was found was denoted as embryonic (E) 0.5. Pregnant dams were euthanized by CO_2_ asphyxiation at E8.5 and embryos were harvested into 1x PBS + 0.01% Bovine Serum Album (BSA) and kept on ice until dissection. We dissected embryos in cold 1X PBS + 0.01% BSA, removing all extra-embryonic tissues. The yolk sac and amnion were cleaned of maternal contaminants and were used for DNA extraction (see below). We transferred dissected embryos immediately to freshly prepared and chilled 4% paraformaldehyde, pH 7.4, overnight at 4°C.

### Whole-mount Immunostaining

After fixing, we washed fixed embryos for 3 × 30 min in 1X PBS + 0.1% Tween20 (PBS-W) at room temperature. We then incubated fixed embryos in Phalloidin Alex Fluor 568 (Invitrogen A12380) overnight at 4°C. We then washed 3 × 30 min in PBS-W at room temperature and blocked for 2 hr in 1X PBS + 1% normal goat serum + 0.5% TritonX-100 + 0.01% NaN_3_ (Blocking Buffer). We then incubated samples with primary antibodies diluted in blocking buffer overnight at 4°C. We washed the embryos 3 × 30 min in PBS-W and then incubated in secondary antibodies diluted in blocking buffer for 2 hr at room temperature. Finally we stained embryos with Hoechst (1:100, ThermoFisher H3570) for 10 min at RT before washing 3 × 30 min in PBS-W at room temperature and storing in 1X PBS + 0.01 % NaN_3_ 4°C until mounting. Antibodies were used at the following concentrations: rabbit anti-non-muscle myosin heavy chain IIB (1: 400, BioLegends 909902) and goat anti-rabbit Alexa Flour 488 (1: 500, ThermoFisher A-11008).

### Whole-mount Imaging

We whole-mounted stained embryos in Aqua-poly/Mount (Fisher Scientific NC9439247) between two #1 glass coverslips. We adhered the coverslips to a glass microscope slide with clear nail polish, such that the apical/dorsal side of the NT faces the microscope objective (Figure 1A). We imaged samples on an inverted Zeiss LSM 900 equipped with a C-Apochromat 40x/1.20 W Korr water objective and pinhole size of 1 airy unit. We acquired images with a 9.6355 pixels/µm XY resolution and Z-spacing of 0.45 µm. We tiled-scanned images so that there was at least a 10% overlap between fields of view.

### Shell Projections

We used FIJI to convert LSM files to multi-image TIFFs as well as Image Sequences made from the Myosin IIB and DNA channels (Figure 1B). We passed image sequences to a trained Ilastik model, which primarily uses intensity to classify pixels as either part of the sample or background. We included the DNA stain to help fill in pores of the more junctional myosin IIB and created a solid mask at the most apical planes. This mask was then refined using a custom MATLAB script that fills in any remaining gaps and smooths out edges. The refined mask and the TIFF were combined to select the 10 most apical pixels (at total thickness of 4.5 µm) at every XY position of the image. The maximum intensity of these pixels was then calculated for each channel to create the final shell projections. For subapical projections used to normalize intensity (see below), we projected the 15^th^ most apical pixel at every XY position of the image.

We combined individual channel projections into composites (Figure 1C). We then manually stitch together composite shell projection in FIJI to reconstruct the brain NT using the Pairwise Stitching plugin (settings: Fusion method – Intensity of random input tile, Check Peaks – 5, Ignore zero values when fusing, display fusion, computer overlap, subpixel accuracy, x=0, y=0, registration channel image 1 and 2 – average all channels)^56^.

### Cell Segmentation

We used the F-actin channel to segment the reconstructed images using Cellpose3 and one-click denoising ^36^(Figure 1D). We trained a custom segmentation model using the pretrained cyto3 model as a base along with a curated training set of our individual field of view projections. We performed manual corrections of segmentations and deleted cells in the surface ectoderm, which had characteristically large apical nuclei and area. We exported a segmentation PNG and cell outlines text file.

### Cell Position Standardization

In each reconstruction, we identified the left and right boundaries of the embryo and the center. We also defined the position of a 100 µm tall band across the midbrain of the embryo and rostral to the midbrain/hindbrain boundary cells. For each side we calculated 100% distance as the distance between the relevant boundary and the center point. We digitally straightened some embryos whose RC was not aligned straight up and down. To calculate the angle of rotation, we drew straight line from the bend in the NT on each side that denotes the midbrain/hindbrain boundary (Figure 2A-C) in FIJI. We then determined the angle between the line and a straight horizontal line. For these embryos, we identified the boundaries, center, and 100 µm band after rotation.

### Cell Shape and Protein Density Measurements

We used the MATLAB *regionprops* function to measure cell shape and intensity measurements. For images that needed to be digitally rotated, we measured cell area, major and minor axis, and F-Actin and Myosin IIB intensity before rotation. We measured cell center position and orientation after rotation. To convert cell X position to percent distance from midline we first calculated the distance between the left and right boundary and the center point. For X-position of all cells, we subtracted the X position, so that cells to the left of the center have a negative position and cells to the right have a positive position. We then divided the X-position of cells on the left with the left-center distance, and likewise with cells on the right with the center-right and multiplied by 100% to rescale positions. Finally, we took the absolute value of the percent distance of cells to pool together cells on the left and right sides of the embryo. We calculated cell aspect ratio as major axis: minor axis. For cell orientation, we flipped the sign of orientation for cells on the left side to pool together cells on the left and right side of the embryo. We then took the absolute value of the orientation so that 0° is parallel to the ML axis and 90° is parallel to the RC axis.

To measure protein density, we calculated the total protein intensity of each cell in the apical projection and divided it by the cell area. As we prepared samples in different batches, due to litter size constraints and embryo age variability within a litter, we sought to normalize intensity between cells. We reasoned that even though the actual intensity may vary from batch to batch, the ratio of intensity at apical and subapical planes should remain constant. Therefore, we calculated the average intensity of the entire subapical projection of an embryo and then divided the protein densities of its cell by that average.

### Live Embryo Culture

We set up timed matings of CD-1 mice for a few hours in the morning and the day a plug was identified was designated Embryonic Day (E) 0. We euthanizes pregnant dams by cervical dislocation at E8 and harvested embryos for whole embryo culture as previously described^57^. We dissected embryos in DMEM + 10% fetal bovine serum (FBS) prewarmed to 37°C, leaving the ectoplacental cone, yolk-sac, and amnion intact. We then transferred embryos to vials containing neat, 0.45 µm-filtered rat serum warmed to 37°C and gassed with 5% O_2_, 5% CO_2_ and 90% N_2_. We placed sealed vials into a roller culture incubator for at least 2 hours to allow embryos to acclimate to culture conditions before laser ablation.

### Live Embryo Laser Ablations

For all ablation procedures, we pre-warmed and maintained all medias, dishes, and incubators at 37°C. We dissected, positioned and ablated embryos individually and sequentially using a modified version of a previously developed method^42^. We monitored embryos for a steady heartbeat throughout our procedures as an indication of health. We started by transferring embryos to a 55 mm petri dish containing a 4% agarose-DMEM base submerged in DMEM + 10% FBS. We breached, but left attached, the yolk sac and amnion to reveal the neural tube and transferred embryos to a 1:100 CellMask Deep Red (Invitrogen C10046) in DMEM + HEPES for 3 min to stain membranes. We transferred embryos back to the petri dish and used microsurgical needles from 11–0 Mersilene (TG140-6; Ethicon) and 10–0 Prolene (BV75-3; Ethicon) to position the cranial neural folds upwards and restrict movement. We kept embryos on a Tokai Hit Thermo Plate III (CellSeed) set to 37°C during mounting.

We performed ablations on a Zeiss Examiner LSM880 confocal microscope using a 20×/NA1.0 Plan Apochromat dipping objective with adjustable refractive index collar set to 1.33 and a SpectraPhysics Mai Tai eHP DeepSee multiphoton laser with a nominal power of 1,828 mW (710 nm wavelength, 100% laser power, 4.1 μs pixel dwell time, 0.21 μs pixel size, one iteration). We ablated a 36 µm line (6x the average diameter of cells in the neural folds) on the apical surface of the lateral neural folds or midline along the rostral-caudal or medial-lateral axis the embryo. Ablations instantly vaporized a thin line of tissue^58^. We imaged every 0.416 sec for a total of 20 frames per ablation, with the ablation occurring between frames 2 and 3. We ablated each embryo up to 5 times within a 15 min window after initial breaching of the yolk and amnion, with each ablation occurring in a non-overlapping field of view. After ablations, we washed embryos in 1X PBS, dissected and collected the yolk and amnion for DNA extraction (see below). We fixed and stained with Phalloidin as described above before storing samples in 1X PBS + 1 % NaN_3_.

### Initial Recoil Velocity Analysis

For each ablation, we extracted frames 2 (pre-ablation) and 3 (post ablation) in FIJI. We corrected for noise using the PureDenoise plugin (settings: Automatic – Global, Cycle-spins = 4, Multiframe = 1). We then brought the images into register using the StackReg plugin^59^, set to Rigid Body. To measure initial recoil velocity, we selected points that were 1) clearly visible in both frames, typically cell vertices, 2) abutted opposite edges of the ablation, and 3) were near the center of the ablation path. Orthogonal control points were selected to be adjacent but parallel to the ablation, and parallel control points were outside the ablation path (Figure 4B). We measured a straight-line distance between pairs of points perpendicular to the ablation path. We repeated this measurement 3 times before and 3 times after ablation. We then subtracted the post and pre ablation distances to get 3 values for change in distance, which we averaged into one final value. We divided this average change in distance by the time between frames to calculate initial recoil velocities.

We performed all measurements blinded to information regarding the region or orientation of the ablation, embryo sex or age, or the ordering in which the ablations were performed. We performed a total of 270 ablations across 83 embryos. We removed recoils for any of the following reasons: The ablation was performed in a region outside of the midbrain midline or lateral folds; We were unable to determine the sex or somite stage of the embryo; Bubbles formed in the ablation path; The background embryo had severe movement artifact that prevented accurate measurements. We defined an embryo to have severe movement if the absolute value of the orthogonal or parallel control recoil was greater than the measured initial recoil velocity. A total of 155 ablations across 71 embryos were ultimately used for analysis in this study.

### DNA Extraction and Embryo Sex Determination

For FVB samples, we boiled yolk and amnions in 50 µl 50 mM NaOH at 95° for 30 min. We neutralized the reaction using 15 50 µl 1M pH 7.4 Tris. For CD1 samples, we digested yolk and amnions in 1 mM Proteinase K in Direct PCR lysis buffer (Viagen 301-C) for 1 hr at 55°C to extract DNA. We inactivated samples for 15 minutes at 95°C before use. From both preparations, we used 1 µl of extract for PCR. We determined sex using forward (5′-CTGAAGCTTTTGGCTTTGAG-3′) and reverse (5′-CCACTGCCAAATTCTTTGG-3′), which target *Jarid1c* and *Jarid1d* genes on the X- and Y-chromosome respectively^60^. The primers anneal at 54°C and produce a 331 bp and 302 bp fragment for the X- and Y-chromosomes respectively when separated on a 2% agarose gel.

### Embryo Somite Staging

Somites begin to appear near the hindbrain/spinal boundary as round cell masses flanking and ventral to the NT. As somites mature, they form a central ring of F-actin and myosin IIB. While it is possible to see some F-actin and myosin IIB enrichment earlier in the formation of somites, we only counted somites if they had an actomyosin ring. We used two methods to visualize and count somites on the embryos. For glass mounted embryos, we imaged samples on an inverted Zeiss LSM 900 equipped with a EC Plan-Neofluar 10x/0.30 M27 air objective and pinhole size of 1 airy unit. We acquired images with a 3.2055 pixels/µm XY resolution and Z-spacing of 4.71 µm. We tiled-scanned images so that there was at least a 10% overlap between fields of view. For laser ablated embryos, we transferred embryos to a petri dish with a 4% agarose in PBS base submerged in 1X PBS and counted the F-Actin enriched somites on a Leica MZFLIII stereomicroscope.

### Statistical Analysis and Figure Assembly

We conducted statistical analyses and graph generation in Python using the Seaborn, Scipy, and Statannotation libraries. We performed an F-test to determine homogeneity of variance between the two groups compared. In instances where the resulting F-test p-value was > 0.05, we performed a t-test assuming equal variance. Otherwise, we conducted t-test assuming unequal variance. We classified values as follows: p-values > 0.05 are nonsignificant (n.s); < 0.05 are shown as “*” on plots; <0.01 were shown as “**” on plots. For fixed image cell shape and protein intensity analysis (Figure 2H-J, Figure 3E-H, Figure 5E-H), N = 5 embryos for all conditions and n, or the number of cells in each 10% is ≥150 cells for each embryo. For live laser ablation dates N is the number of cuts performed in the indicated region or location and are listed from left to right for the following figures: 4C 155, 155, 155; 4E – 100, 55; 4G – 74, 34, 26, 21; 5J – 14, 20, 11, 10; 5L – 30, 44, 10, 16. Neural tube cartoons were sketched in Adobe Illustrator and colored in Adobe Photoshop. All other graphical components were created in Adobe Illustrator. Complete figure panels were assembled in Adobe InDesign.

## ACKNOWLEDGEMENTS

We would like to thank members of the Martin, Galea, Copp, and Greene groups for their feedback and support throughout the course of this work. We thank Dr. Ann Sutherland for initial training in live mouse culture, Dr. Rosie Marshal for her support at UCL, and Dr. Maria Garcia-Garcia for initial introduction into mouse husbandry. We thank Dr. Andy Copp for his helpful conversations about this work, Thomas Evans for additional commentary and editing on this manuscript during writing, and Dr. Michael J. Redlich for his continued encouragement and support with editing. This work was supported by: the National Science Foundation Graduate Research Fellowship under Grant No. 2141064 (J.D.); the National Institute of General Medical Sciences of the National Institutes of Health F31 GM154461 (J.D.) and R35GM144115 (A.C.M); the UK Medical Research Council MR/W019876/1 (G.L.G); the Royal Society RG\R2\232082 (G.L.G); and the Company of Biologists Travel Fellow DEVTF240051496 (J.D.)

## SUPPLEMENTAL FIGURES

**Figure 3 – Supplement 1:**
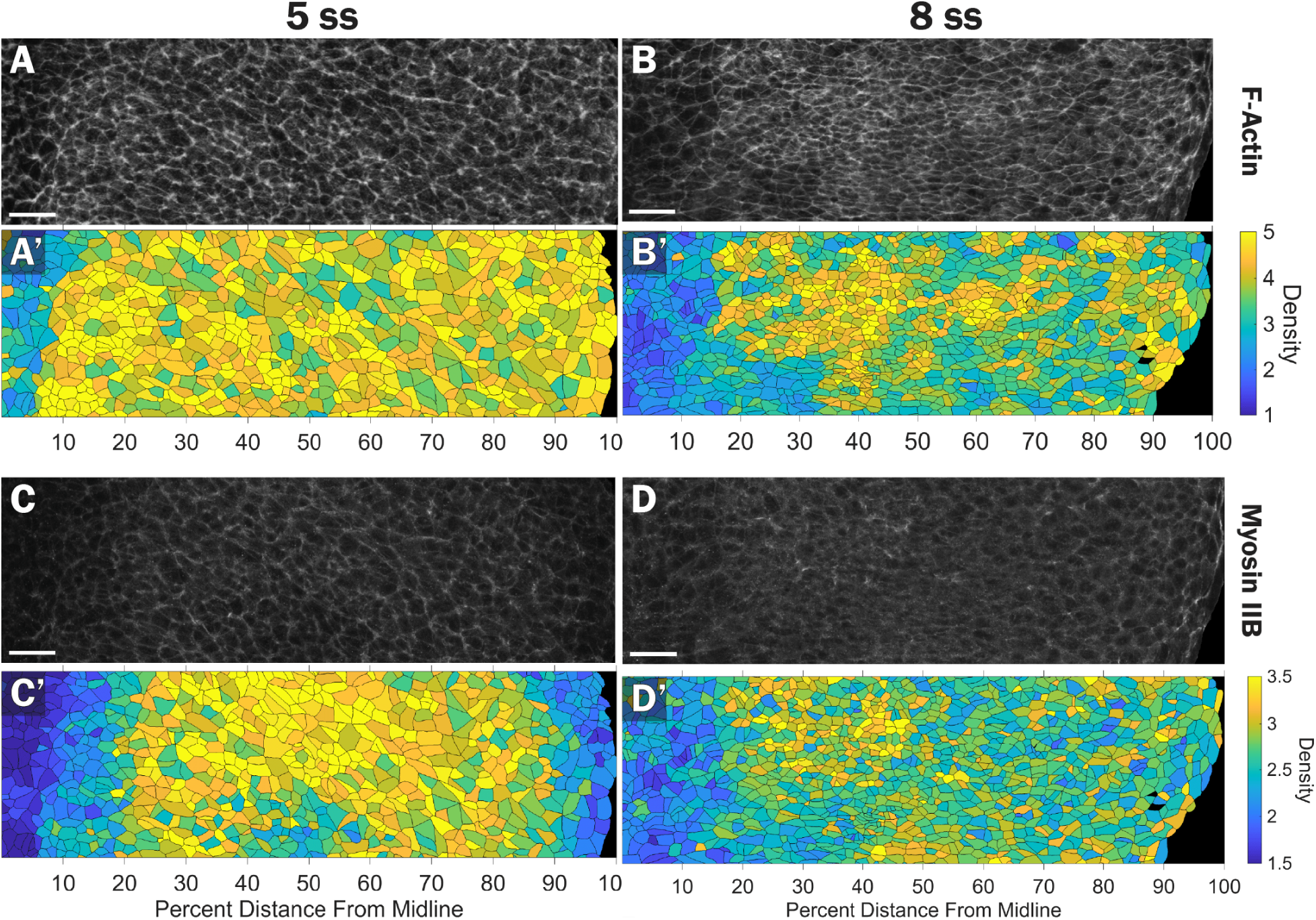
The male midbrain also exhibits a midline-lateral gradient of actomyosin. **A-D)** Representative images of female midbrains stained for A-B) F-actin, and C-D) Myosin IIB at A, C) 5 and B, D) 8 ss. **A’-D’**) Segmented images color coded by density of protein expression from low (dark blue) to high (yellow).

**Figure 4 – Supplement 1:**
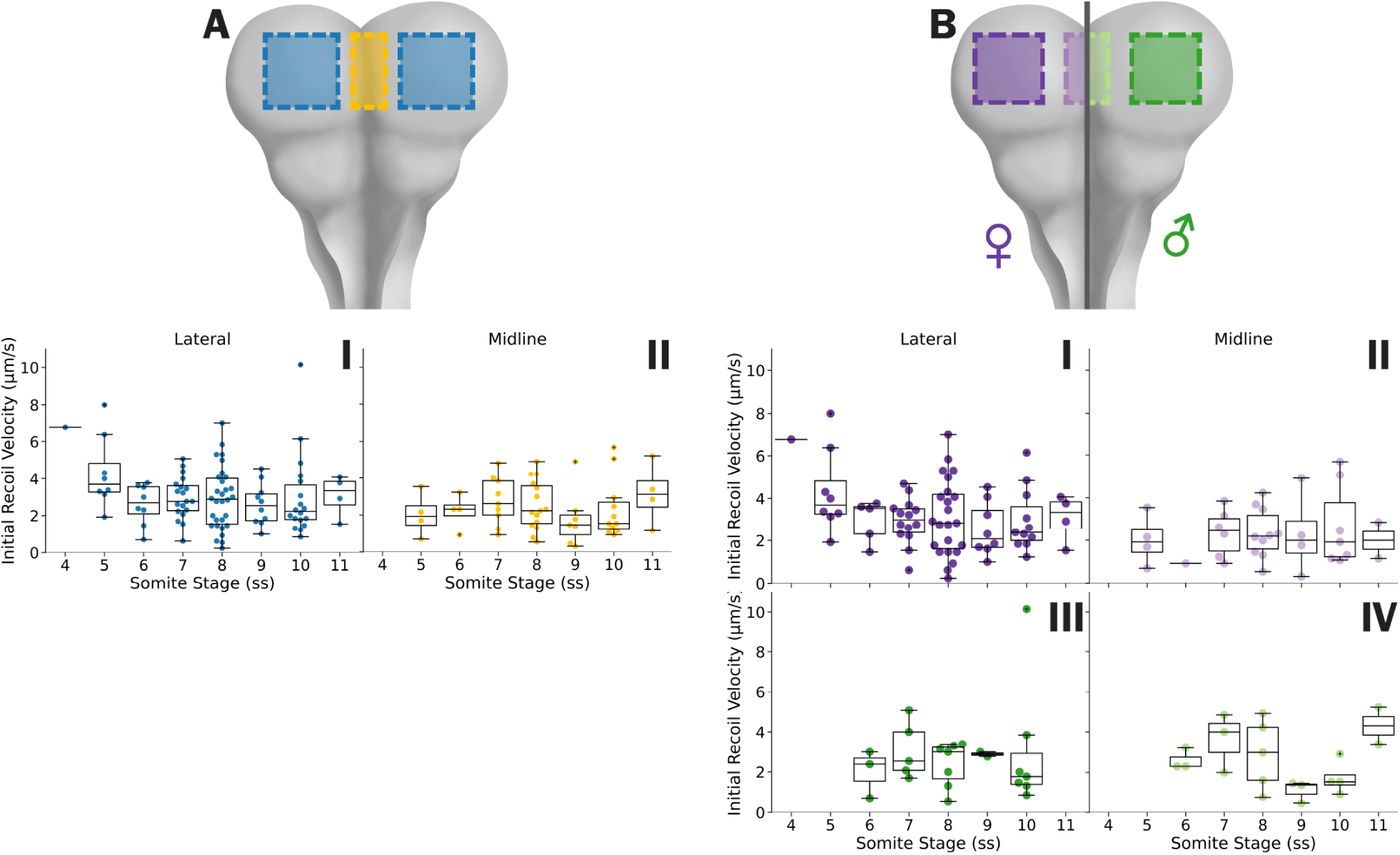
Initial recoil velocities show no difference over developmental time. **A-B)** A cartoon schematic of location where initial recoil velocities were measured for their respective subplots (**I-IV**). A and B) Shaded areas indicate regions where initial recoil where measured. In B) female measurements are in purple hues and males are in green hues. In I-IV) box plots show the median and interquartile range.

**Figure 5 – Supplement 1:**
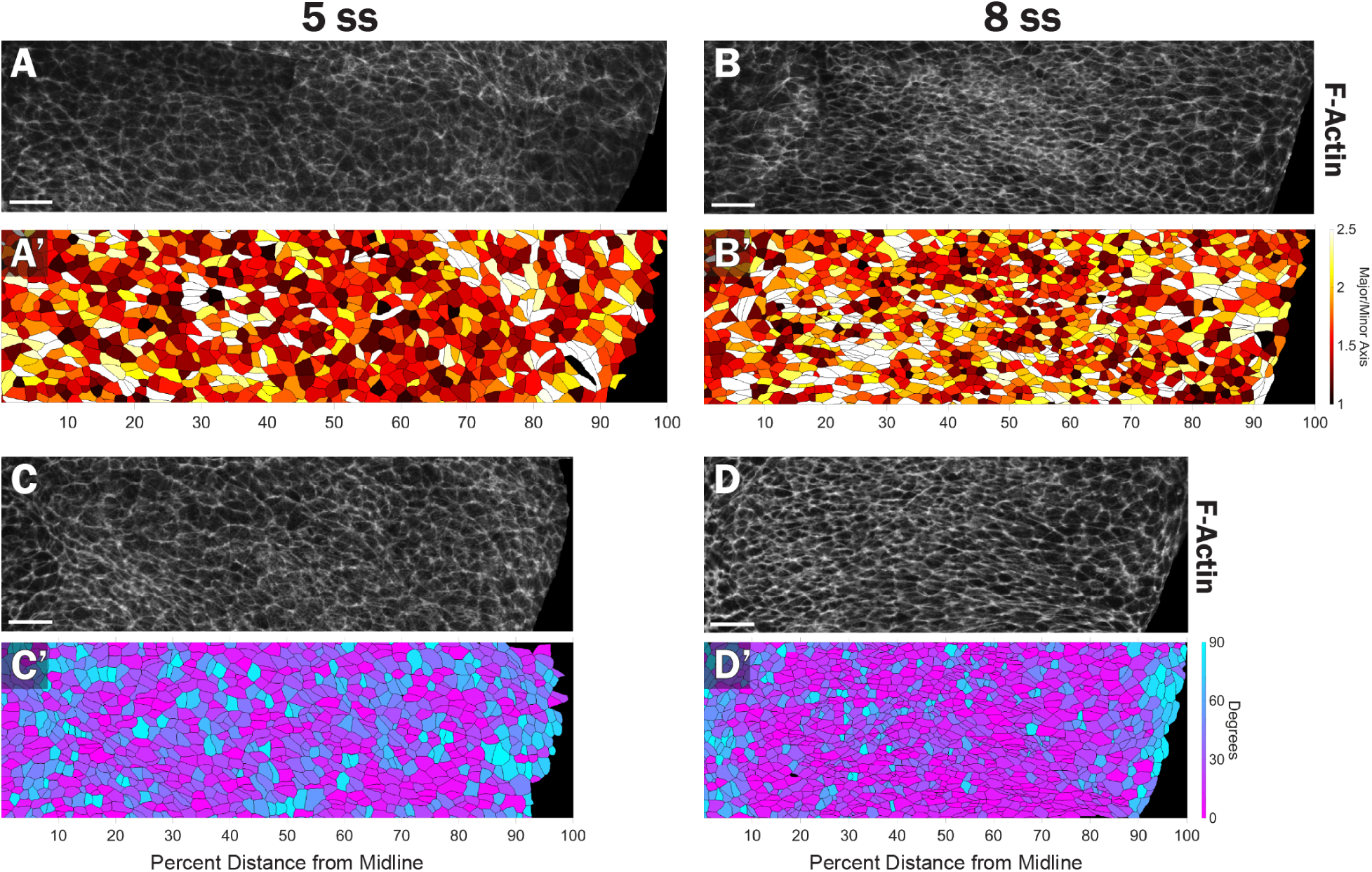
The lateral neural folds of the male midbrain exhibit a cell orientation bias. **A-D)** Representative F-actin images of male midbrains at A, C) 5 and B, D) 8 ss. A’-B’) Segmented images color coded by aspect ratio from uniformly round (black) to elongated (white). **C’-D’**) Segmented images color coded by degree from midline, with 0 (pink) being parallel to the midline-lateral axis and 90 (blue) being parallel to the rostral-caudal axis.

**Figure 5 – Supplement 2:**
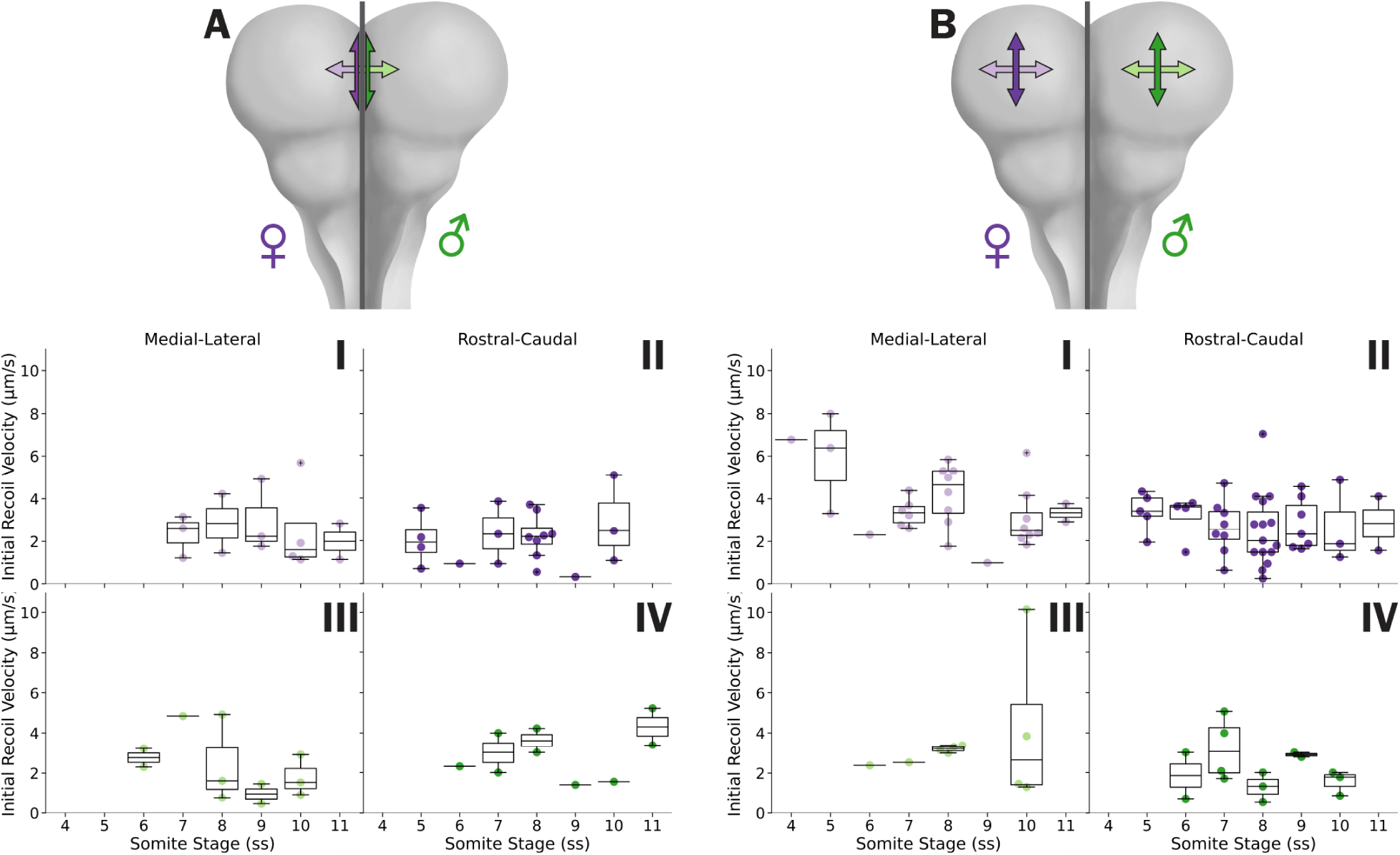
Initial recoil velocities show no difference over developmental time when split by direction. **A** and **B)** A cartoon schematic of location and direction where initial recoil velocities were measured for their respective subplots (**I-IV**). Double headed arrows indicate the direction inferred tension was measured in the A) midline and B) lateral neural folds. In I-IV) box plots show the median and interquartile range. Female measurements are in purple hues and males are in green hues.

## Notes

### Competing Interest Statement

The authors have declared no competing interest.

### Summary of Updates

Added 4 additional data figures showing patterns of actomyosin density and cell shape across the developing neural tube.

